# Spatial Variability in Soil Necrobiome Communities has a Negligible Effect on Postmortem Interval Estimation

**DOI:** 10.64898/2026.07.07.737041

**Authors:** Lesley Hewett, Cydney Rimok, Karen A. Thompson, Shari L. Forbes, Aaron B. A. Shafer

## Abstract

Microbial succession can be used to estimate the postmortem interval (PMI); however, the impact of spatial variability within the cadaver decomposition island (CDI) is not well understood. This study examined spatial variation in necrobiome communities where soil samples were collected over time and across spatial locations from the CDIs of two human body donors. Microbial communities were characterized using 16S rRNA sequencing and statistical modelling of variation and PMI were conducted. Necrobiome community metrics showed no significant differences across anatomical sampling sites within the CDI at a single timepoint. Temporal modelling identified 11 taxa with significant relationships to PMI in one donor, with spatial sampling having a minimal impact on the PMI relationships. Non-linear approaches also identified taxa with likely PMI signals in the second donor. These findings demonstrate that opportunistic sampling can capture robust linear and non-linear PMI signals in later decomposition stages.

## Introduction

When vertebrate decomposition occurs in a terrestrial environment, cadaveric materials enter the surrounding soil, creating a biologically and chemically distinct microenvironment termed the Cadaver Decomposition Island (CDI) [1]. The resulting CDI is a visually distinct area surrounding the remains, characterised by elevated nutrient levels, including carbon, nitrogen, phosphorous and sulfur, altered soil pH, increased electrical conductivity and reduced oxygen levels [1–7]. The pulse of organic material also flushes the host microbiome into the environment, which coalesces with scavenger-derived microbes and indigenous soil bacteria to form the necrobiome, a community of microbial decomposers that are associated with the organism’s remains [8]. The necrobiome community shifts in response to the physicochemical conditions created by decomposition, producing patterns which are predictable and can be modelled to predict time since death, or postmortem interval (PMI) [1,2,9–19]. Although temporal patterns in microbial relative abundance within the CDI are documented, there has been relatively little research aimed at understanding how the necrobiome composition varies within the CDI.

A growing body of research suggests that CDIs are spatially heterogenous systems, with differences arising as a result of inherent variability in human anatomy and decomposition processes. Distinct microbial communities are associated with different body locations, which are dispersed into the surrounding soils in a spatially variable way due to differential decomposition between regions, anatomically focused insect activity, and uneven release and migration of decomposition fluids [14,15,20–23]. The inherent asymmetry of these processes creates spatial variation in soil conditions, with nutrients, isotopes, lipids and other decomposition products displaying analyte-specific distributions within the visible extent of the CDI [3,24–27]. Because microbial community composition and structure are fundamentally shaped by these soil conditions, variation of these properties within a CDI indicates that necrobiome distribution may not be spatially uniform [5,13,15,28,29]. Combined, this evidence suggests that CDIs are not chemically or microbially homogeneous environments, highlighting important considerations for necrobiome sampling strategies and subsequent PMI estimates.

PMI prediction models are typically built using microbial abundance data generated from a timeseries of samples that are intended to capture a representative subset of the necrobiome population at each sampling point while also minimizing the disturbance this causes to the decomposition environment [30,31]. In other disciplines studying soil microbes, the impact of spatial variability within a study area is mitigated by collecting multiple subsamples at each time point and combining them into one representative composite sample [32–34]. Several necrobiome studies have adopted this approach, often collecting subsamples from below the torso of the remains; however, lifting the cadaver for repeated sample collection impacts the carcass temperature and may disrupt insect colonization, two factors known to strongly influence the necrobiome and decomposition progression [35,36]. To reduce disturbance, other approaches collected samples from the area immediately surrounding the remains or restrict sampling to locations proximal to specific anatomic sites. These approaches minimize manipulation of the remains, but are confined to a smaller sampling area, which can be limiting when the study requires frequent sampling or covers an extended postmortem interval. In either approach, repeatedly removing soil from the system may introduce local aeration, which can impact anaerobic decomposition and microbial community composition [3,4,37]. As a result, existing necrobiome studies use a range of sampling strategies that reflect different approaches to balance the desire to collect representative samples with the need to minimize the impact on the decomposition environment. The variety of sampling strategies used in temporal necrobiome studies suggests that there is ongoing uncertainty about spatial variability within the CDI, highlighting a need to better understand differences in the necrobiome within the CDI at a given time point.

To explore these questions, we analyzed soil samples collected from the CDI of two human body donors decomposing at the Research in Experimental and Social Thanatology (REST) human taphonomic facility. We hypothesized that microbial communities within the CDI would exhibit spatial heterogeneity at a given time point during human decomposition, and predicted that necrobiome community composition would differ among soil samples collected from distinct anatomical regions within the CDI [13,15,18]. Further, we hypothesized that spatial variation could influence the detection of temporal relationships between microbial abundance and PMI. We predicted that informative relationships between necrobiome composition and PMI would be detected, despite spatial heterogeneity within the CDI [17].

## Methods

### Experimental Setup

Samples were collected from two deceased individuals (donors) at the Research in Experimental and Social Thanatology (REST) human taphonomic facility, located on a 1600 m^2^ lot in Bécancour, Québec, Canada [38]. All individuals provided consent to participate in the willed body donation program through the Université of Québec at Trois-Rivières (UQTR). Bécancour is located within the Dfb climatic zone as defined in the Koppen-Geiger classification system, which is characterized by a cold climate, with no significant precipitation difference between seasons, and a warm summer [39]. REST is situated in a mixed temperate forest dominated by maple and spruce trees, and sandy loam soil and experiences seasonal temperatures ranging from -40°C in the winter to 40°C in the summer [38,40]. Meteorological data is recorded by an on-site weather station, which collects data on temperature, humidity, rainfall, solar radiation, wind speed, and direction [38]. Maximum and minimum daily temperatures were obtained from the onsite weather station, with any missing data supplemented using records from the nearest Environment and Climate Change Canada weather station.

The two donors were placed on the surface of the ground, unclothed in a supine position, having been refrigerated at 4°C between death and placement (within 3 days of death). Donors were placed on the soil surface under anti-scavenging cages, which allow insect access, but prevents vertebrate scavenging.

Cages were spaced at least 2.3m away from each other, avoiding all known sites previously used for body deposition. The decomposition stage of each donor was assessed qualitatively using visual characteristics of the remains and CDI, as described by Payne [41] and Carter et al [1]. Relevant information about the donors in this study is outlined in Table 1. The identification number for each donor was maintained based on the REST identification system (in this case, Donor 2 and 21).

**Table 1.** Donor Information Summary.

| Metric | Donor 2 | Donor 21 |
| --- | --- | --- |
| Age at death (years) | 71 | 84 |
| Sex | Male | Male |
| Body mass (kg) | 70 | 68 |
| Height (m) | 1.73 | 1.72 |
| Body Mass Index (BMI) | 23.4 | 23 |
| Date of death | August 7, 2020 | November 26, 2022 |
| Date of placement at REST site | August 10, 2020 | November 28, 2022 |
| Cause of death | Hepatic cirrhosis, anuric renal insufficiency | Sigmoid colon perforation, diverticulosis |
| Postmortem Interval Study Range | 1000 - 1186 days | 159 – 345 days |
| Accumulated Degree Day Study Range | 8255 - 11284 ADD | 414 - 3443 ADD |
| Decomposition Stage During Study | Dry | Advanced Decay |

### Sample Collection and Preparation

Soil sample collection for this study began in May 2023 and continued until late fall of that year, covering a span of 186 days. Throughout this period, Donor 2 was in the dry remains stage of decomposition, and presented as exposed bone with sparse desiccated tissues and a visually uniform, dry CDI showing plant regrowth. Donor 21 was in Advanced Decay during the sampling period with liquified, sunken soft tissue, bone exposure, decomposition fluid accumulation around the trunk, and a defined, moist CDI lacking any vegetation. While some of these conditions progressed throughout sampling, Donor 21 did not reach the dry remains stage within the study period.

During this time, single soil cores were collected from any CDI location within 15 cm of the border of the torso of D2 and D21, at intervals ranging from 13 to 33 days between collection. These samples were used for the temporal analysis. Approximately halfway through the temporal sampling period (PMI 1098 for Donor 2 and PMI 257 for Donor 21), six soil cores were collected from each donor at specific CDI sites: shoulder, pelvis, and knee on both the left and right sides of the body (Figure 1). These samples were used for spatial (CDI) comparisons at this single timepoint and incorporated into the temporal analysis.

**Figure 1.**
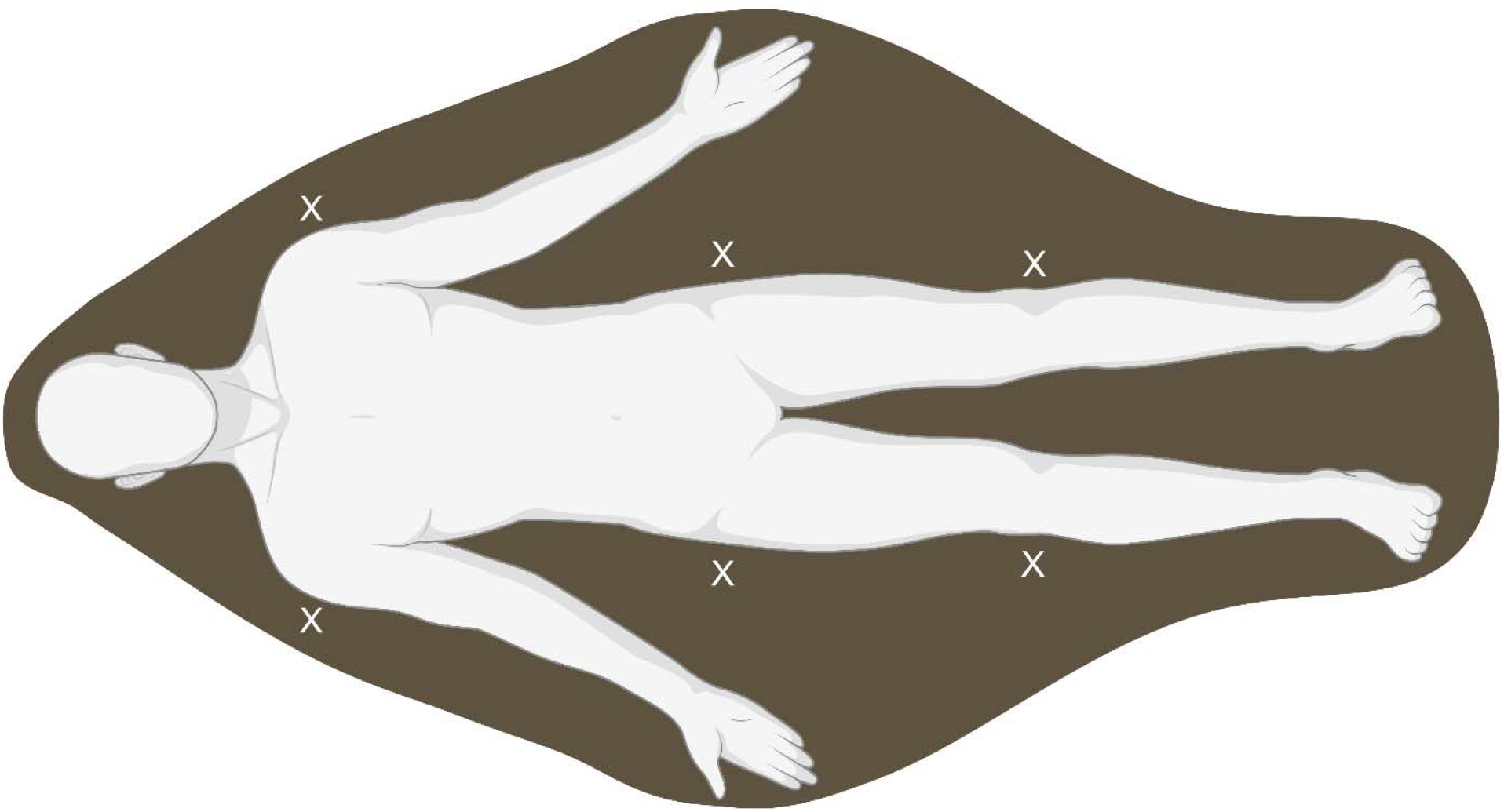
Visual representation of the cadaver decomposition island (dark area) surrounding decomposing human remains. Locations marked with an ‘X’ represent the areas where spatial samples were collected. Temporal samples were collected from any location within the CDI between the shoulders and knees. Image created using BioRender.

At the time of sample collection, control samples were also collected from a site inside the facility but at least 3.8m away from any known deposition site as well as a site ∼18m outside of the facility entrance, to allow for monitoring of site-specific community variation unrelated to decomposition. Soil cores were collected using a sterilized 2.5 cm-diameter soil corer to a depth of 5 cm and transferred into a sterile whirl-a-pack. Cores were manually homogenized before being stored at -20°C, where they were held until analysis.

DNA was extracted using the DNeasy PowerMax Soil Kit following the manufacturer’s protocol (Qiagen). Library preparation followed the Earth Microbiome 16S Illumina Amplicon Protocol V.2 [42]. Briefly, this involved amplifying the V4 region of the 16S ribosomal RNA gene (16S rRNA) using the 515F (Parada) and 806R (Appril) primer pair [43–45]. Barcoded amplicons were pooled in equimolar amounts and sequencing was performed in 250 base pair paired end reads on an Illumina MiSeq sequencer at The Centre for Applied Genomics (Toronto, Ontario, Canada).

### Data Analysis

#### Sequence Processing and Bacterial Community Analysis

Raw paired-end FASTQ files were imported into QIIME2 (v.2024.2) [46] and primer sequences were removed using Cutadapt (v.4.6) [47] with no indels allowed. Reads were merged with VSEARCH (v.2.27.0) [48] and quality filtered using the q-score method via the q2-quality-filter plugin [46].

Denoising, dereplication, and chimera removal were performed with Deblur (v.1.3.0) [49], using a trim length of 250 bp. Rare features occurring in <0.0005% of reads in the total dataset were removed, as recommended for Illumina generated data [50]. Taxonomic classification was performed using the classify-sklearn method [51] and a pre-trained SILVA 138 classifier [52]. ASVs unclassified at the phylum level or assigned as mitochondria or chloroplast were removed.

Count tables were exported from QIIME2 into RStudio (v.4.4.1) [53] for analysis and visualization. The dataset was subset to include only spatial or CDI samples and associated control soils, ASVs were merged at the phylum level using the tax_glom function of the phyloseq (v.1.48.0) [54] package and relative abundance was calculated for each phylum. The 12 phyla with the highest mean relative abundance were used to generate a stacked bar plot to visualize differences in community membership between spatial samples. Differences in relative abundance between the left and right sides of the spatial samples were assessed for each donor using the Wilcoxon rank-sum test. The p-values for each taxa were adjusted using the Benjamini–Hochberg (BH) method, to account for multiple significance testing [55]. The phyloseq (v.1.48.0) [54] package was used to rarefy samples in this subset to a depth of 3455 sequences, which retained all relevant CDI samples. Rarefied ASVs were used to calculate alpha diversity metrics, including the Shannon Index and the number of observed features, which were evaluated to assess within-sample diversity in the spatial samples. Values for each donor and control soils were visualized using ggplot2 (v.3.5.1) [56]. Differences in alpha diversity values between the left and right sides of the spatial samples were assessed for each donor using the Wilcoxon rank-sum test and BH-adjusted p-values.

Beta diversity was assessed using Bray–Curtis dissimilarity distances and visualized with a Principal Coordinates Analysis (PCoA) using the plot_ordination function of the phyloseq (v.1.48.0) [54] package. Differences in microbial community structure between spatial groups (e.g. left-to-right) were evaluated using the adonis2 function of the vegan package (v. 2.6-6.1) [57] and tested using a PERMANOVA, while within-group dispersion was assessed using the betadisper function of the vegan package (v.2.6-6.1) [57] and tested using an ANOVA. Environmental fitting was performed using the envfit function of the vegan package (v.2.6-6.1) [57] to identify taxa whose relative abundances were significantly correlated with the ordination axes of the Bray-Curtis PCoAs. Vectors of significant phyla after BH-adjustment were overlaid on the Bray-Curtis PCoA plot to visualize taxa significantly associated with observed spatial patterns.

#### PMI Estimation

Accumulated Degree Days (ADD) are units of time that account for heat exposure above a certain threshold and are typically related to progression of a biological process [23]. ADD were calculated according to the methods in Megyesi et al [23] by summing average daily temperatures from the date of body placement at REST to the date of sampling, using a base temperature of 0°C. Temperature data were collected from an on-site weather station at the REST facility and missing values were assembled from the closest Environment Canada and Climate Change weather station.

In order to capture a range of levels of taxonomic resolution, rarefied ASVs for the entire temporal and CDI dataset were merged at phylum, order and genus levels using the tax_glom function of the phyloseq (v.1.48.0) [54] package. For each taxonomic level, taxa abundance was evaluated at each time point for each donor, and taxa with abundance = 0 in three or more timeseries samples or two or more spatial variability samples were removed from analysis. Abundance counts in the control samples were plotted against time (ADD) as a linear model, and any taxa with statistically significant time signals (p < 0.05) in the controls were removed from the analysis to ensure the taxa being investigated were related to decomposition and not environmental factors. Redundant taxa were removed by identifying higher-rank taxa with abundances identical to a single, lower-rank taxa and retaining the lower rank taxa.

For the remaining taxa, a base linear model was generated using abundance counts plotted against time (ADD) in all timeseries samples (excluding the CDI samples) to identify those with statistically significant temporal signals in the absence of within-timepoint spatial variation. To evaluate the sensitivity of these models to within-timepoint spatial variation (i.e. the CDI), each spatial variability sample (n = 6) was sequentially added to the base model. This created six linear models per significant taxa (one for each sampling location in the CDI) and the resulting slope, R^2^ values and p-values were analyzed to evaluate the effect CDI sampling had on model performance and temporal significance. For each taxa, the p-values for the spatial variability models were adjusted using the Benjamini–Hochberg (BH) method, to account for multiple significance testing [55].

#### ASV Appearance/Disappearance with PMI

Lastly, rarefied ASVs in donor samples were screened to identify taxa that showed zero abundance at two or more consecutive time points at the start of the sampling period (identified as ‘appearance’) or at the end of the sampling period (identified as ‘disappearance’). Taxa that showed the same zero-abundance patterns in the donor samples and control soils were excluded, as were taxa with non-consecutive zero-abundance timepoints and those which had ≥6 zero-abundance time points, regardless of the patterning. For the remaining taxa, log(Abundance) was plotted against sampling date for control soils as well as CDI and temporal samples for both donors to evaluate patterns in the appearance/disappearance of taxa related to PMI including the CDI. Taxa labeled as ‘uncultured’ or ‘unclassified’ at a given rank were reported at the next highest confidently resolved taxonomic level.

## Results

### Bacterial Community Analysis

A total of ∼7.1 million paired-end reads were generated from the 47 soil samples, including 12 spatial variability samples, 17 timeseries samples and 18 control soils. After quality filtering, ∼2.26 million high quality sequences remained representing 3455 unique amplicon sequence variants (ASVs). Two control soil samples were excluded from the analysis as a result of low sequence counts and one timeseries sample was removed since it was part of another study, leaving 44 samples for spatial and temporal analysis.

### Relative Spatial Abundance

Spatial samples and associated control soils were dominated by Proteobacteria, with replicates taken at the control site showing similar compositions and distribution (Figure 2A). The relative abundances of the spatial samples for Donor 2 showed similar communities with some variation. Specifically, Acidobacteriota showed higher proportions in samples from the right side (∼15%) of the CDI than the left (∼4%) (Figure 2A, Supplementary Table 1). A longitudinal gradient of increasing abundance of Proteobacteria from the shoulder to the knee on both sides of the body (+25%) was also observed and was accompanied by patterns of decreasing abundance along the same plane for Firmicutes (-9%), Actinobacteriota (-8%) and Bacteroidota (-4%) (Figure 2A, Supplementary Table 1). Spatial samples for Donor 21 had ∼20% greater proportion of Firmicutes in both pelvic samples, corresponding with sharp reductions (77% - 100%) in most other phyla compared to other sampling sites (Figure 2A, Supplementary Table 1). Wilcoxon rank-sum tests revealed no significant differences in relative abundance values between left and right sides of the CDI of either donor for any taxa (BH adjusted p-values > 0.05).

**Figure 2.**
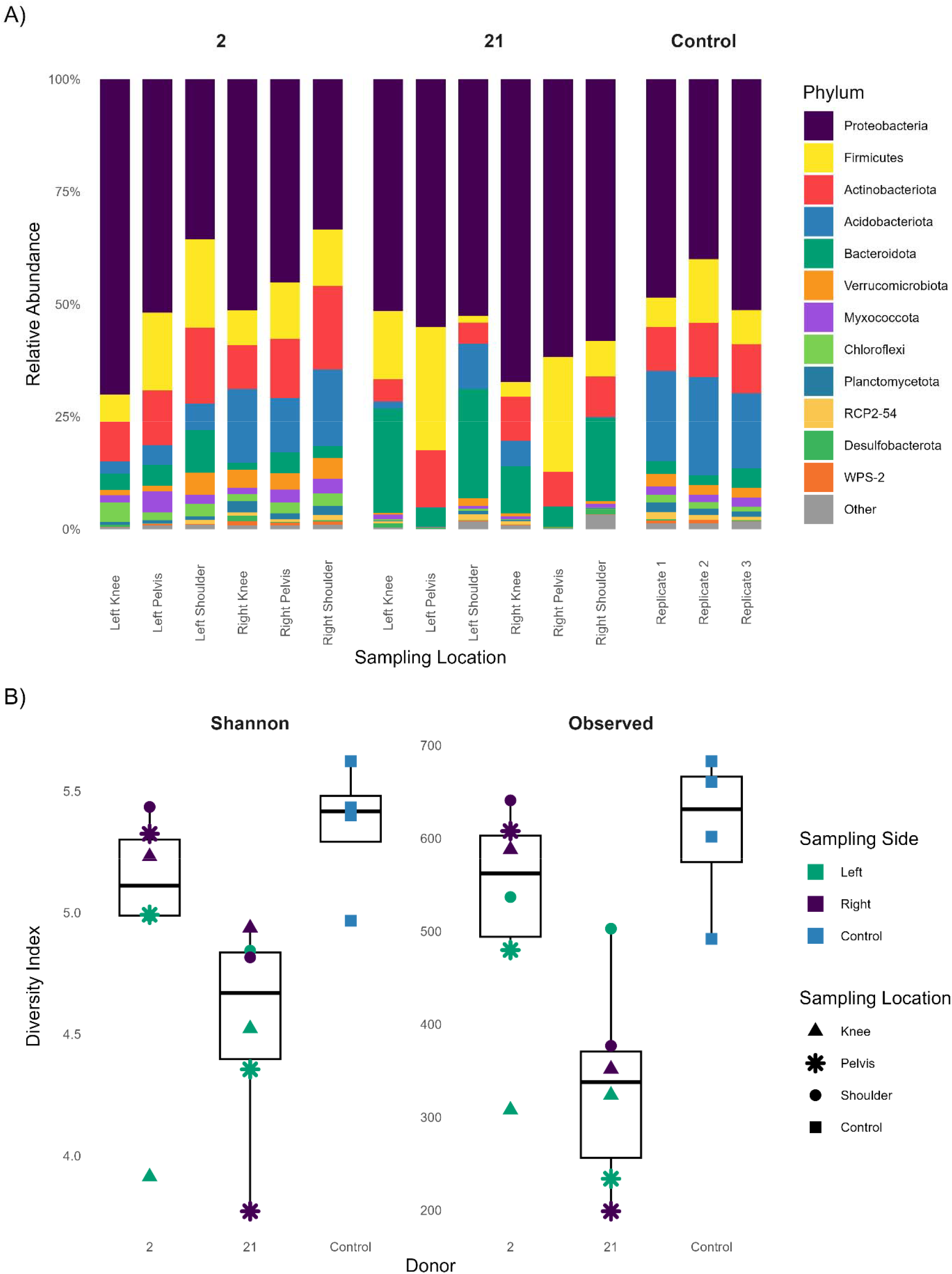
Bacterial Community Comparison Data **A-** Relative abundances of the 12 most abundant bacterial phyla present in the spatial samples for Donors 2 and 21 and the associated control samples. Phyla with abundances ranked 13 and below are grouped together and represented as ‘Other’. Bars are grouped by donor and sampling locations (left and right knees, pelvises and shoulders) with control samples shown as replicates. **B -** Alpha diversity of microbial communities by donor and sampling side. Boxplots show the median (horizontal line), interquartile range (box), and range (whiskers) of alpha diversity metrics (Shannon and Observed Features) calculated from rarefied samples collected from Donors 2 and 21, as well as control soils. Samples are grouped by donor and colored by sampling side, with symbols indicating sampling location.

### Alpha and Beta Spatial Diversity

Alpha diversity metrics (Shannon Index and Observed Features) varied by donor, sampling side and sampling location (Figure 2B). Control soils exhibited the highest diversity and lowest variation in both metrics. Samples from the right side of the CDI from Donor 2 showed higher diversity values similar to the controls for all metrics; however, the values from the left side were consistently lower and more variable (Figure 2B). Samples from the CDI of Donor 21 exhibited the lowest alpha diversity, with communities from the pelvic region having the lowest number of observed features and Shannon Index values of all samples measured (Figure 2B). Despite this, CDI samples from Donor 21 showed more consistency between the left and right sides, with overlapping ranges for both metrics and comparable spread between data points (Figure 2B). Wilcoxon rank-sum tests revealed no significant differences in alpha diversity values between left and right sides of the CDI for either donor or metric (BH adjusted p-values > 0.05). PERMANOVA tests of the Bray Curtis dissimilarity distances showed no significant differences in community composition between sides of the CDI (p >0.05) for either donor (Supplementary Figure 1). Environmental fitting did not identify any taxa that significantly correlated with the ordination for either donor after correcting for multiple testing. ANOVA tests showed no significant differences in within-group dispersion between the left and right sides of the CDI (p>0.05) for either donor.

### PMI Estimation

Linear modelling of the timeseries samples identified 11 taxa with statistically significant linear relationships to ADD (Table 2), with only samples from Donor 2 found to have significant linear relationships with ADD. The addition of the spatial samples (i.e. random CDI location) did not change the overall direction or magnitude of the slope of any model (Table 2). Of note, five of these relationships maintained statistical significance (BH adjusted p-values < 0.05) when varying the spatial sample in the base model (Table 2), meaning they were not impacted by CDI sampling location. Location and taxa that impacted the model significance (p > 0.05) are shown in Figure 3. We identified four ASVs that appeared or disappeared during decomposition, with this pattern not seen (or marginally detected) in the other donor or control samples (Figure 4).

**Table 2.** Summary of model statistics for taxa that showed statistically significant linear relationships with ADD in the base timeseries models. Donor 21 is not included since only models generated using samples from Donor 2 showed a linear relationship with ADD. Taxa listed in italics are at the genus level, all others are at the order level. The statistics associated with each base model are reported in the Slope, R^2^, and p-value columns, and are followed in brackets by the range of values for the same metric from the six spatial models. The models that maintained statistical significance after the addition of all spatial samples are denoted by a (*) in the Spatial Significance column (p-values < 0.05).

| Taxa | Slope | $R^2$ | p-value | Spatial Significance |
| --- | --- | --- | --- | --- |
| Polyangiaceae | -0.02 [-0.02 - -0.02] | 0.67 [0.46 - 0.66] | 0.01 [0.02 - 0.04] | * |
| Micrococcales | -0.02 [-0.02 - -0.02] | 0.56 [0.48 - 0.56] | 0.03 [0.04 - 0.04] | * |
| Burkholderiales | 0.06 [0.06 - 0.06] | 0.55 [0.47 - 0.56] | 0.03 [0.04 - 0.04] | * |
| *WD260* | 0.01 [0.01 - 0.02] | 0.88 [0.14 - 0.88] | 0.001 [0.001 - 0.33] |  |
| *Rhodococcus* | -0.02 [-0.02 - -0.01] | 0.80 [0.02 - 0.80] | 0.003 [0.003 - 0.71] |  |
| *Reyranella* | 0.01 [0.009 - 0.01] | 0.51 [0.40 - 0.50] | 0.05 [0.06 - 0.07] |  |
| *Puia* | 0.04 [0.04 - 0.04] | 0.60 [0.55 - 0.60] | 0.02 [0.02 - 0.02] | * |
| *JG30-KF-CM45* | -0.001 [-0.001 - -0.001] | 0.59 [0.08 - 0.59] | 0.03 [0.03 - 0.47] |  |
| *Gemmatimonas* | -0.002 [-0.002 - -0.002] | 0.60 [0.30 - 0.60] | 0.02 [0.03 - 0.13] |  |
| *Acidothermus* | -0.03 [-0.04 - -0.03] | 0.59 [0.38 - 0.59] | 0.03 [0.05 - 0.08] |  |
| *Acidicapsa* | 0.007 [0.007 - 0.007] | 0.73 [0.64 - 0.72] | 0.007 [0.01 - 0.01] | * |

**Figure 3.**
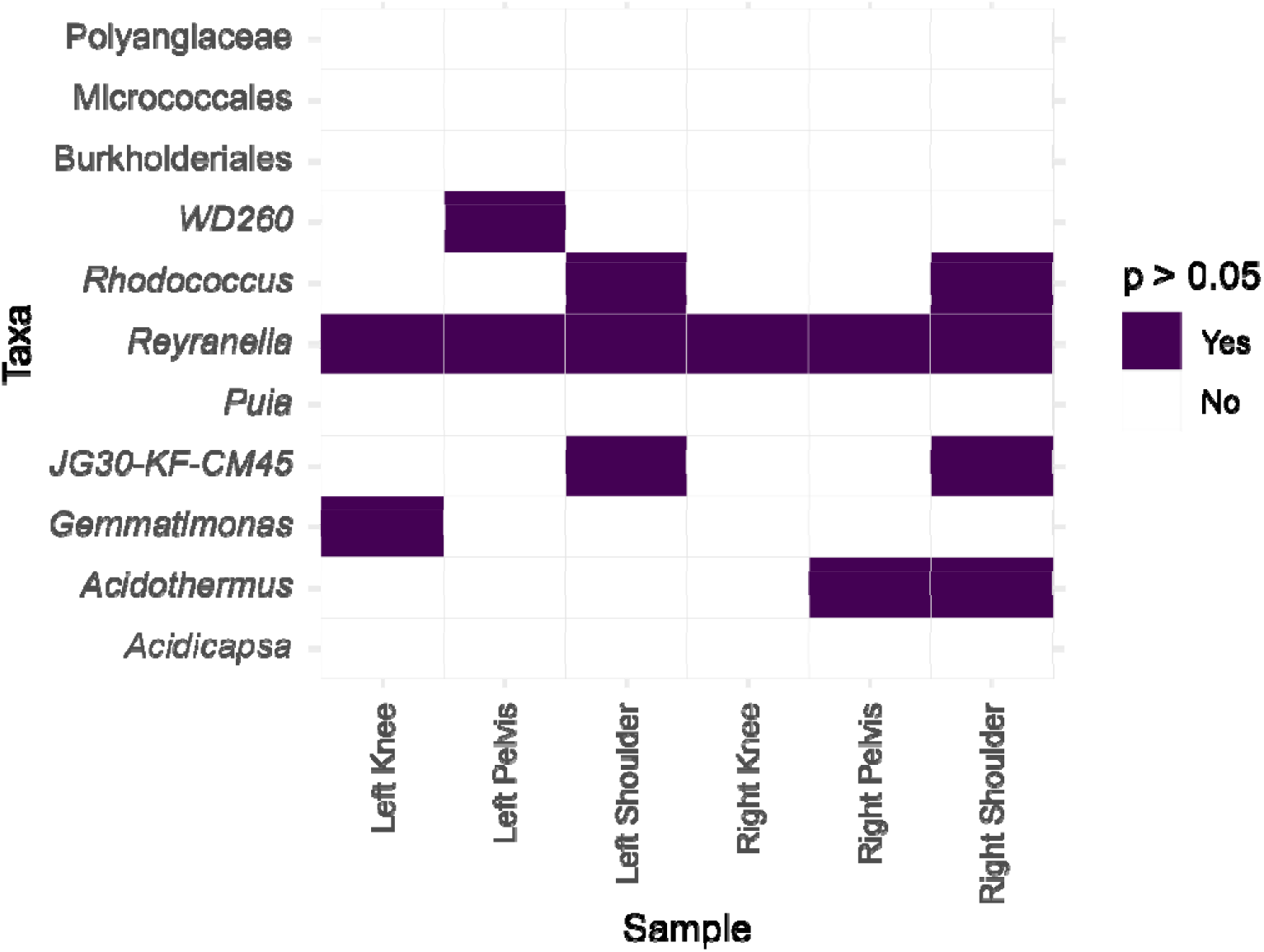
Changes in model significance by taxa and spatial variability sample. Each tile represents a taxon-sample combination, with purple indicating that the addition of that spatial sample to the base model caused the relationship to become non-significant. This highlights the taxa and sampling locations that are most robust to spatial variability.

**Figure 4.**
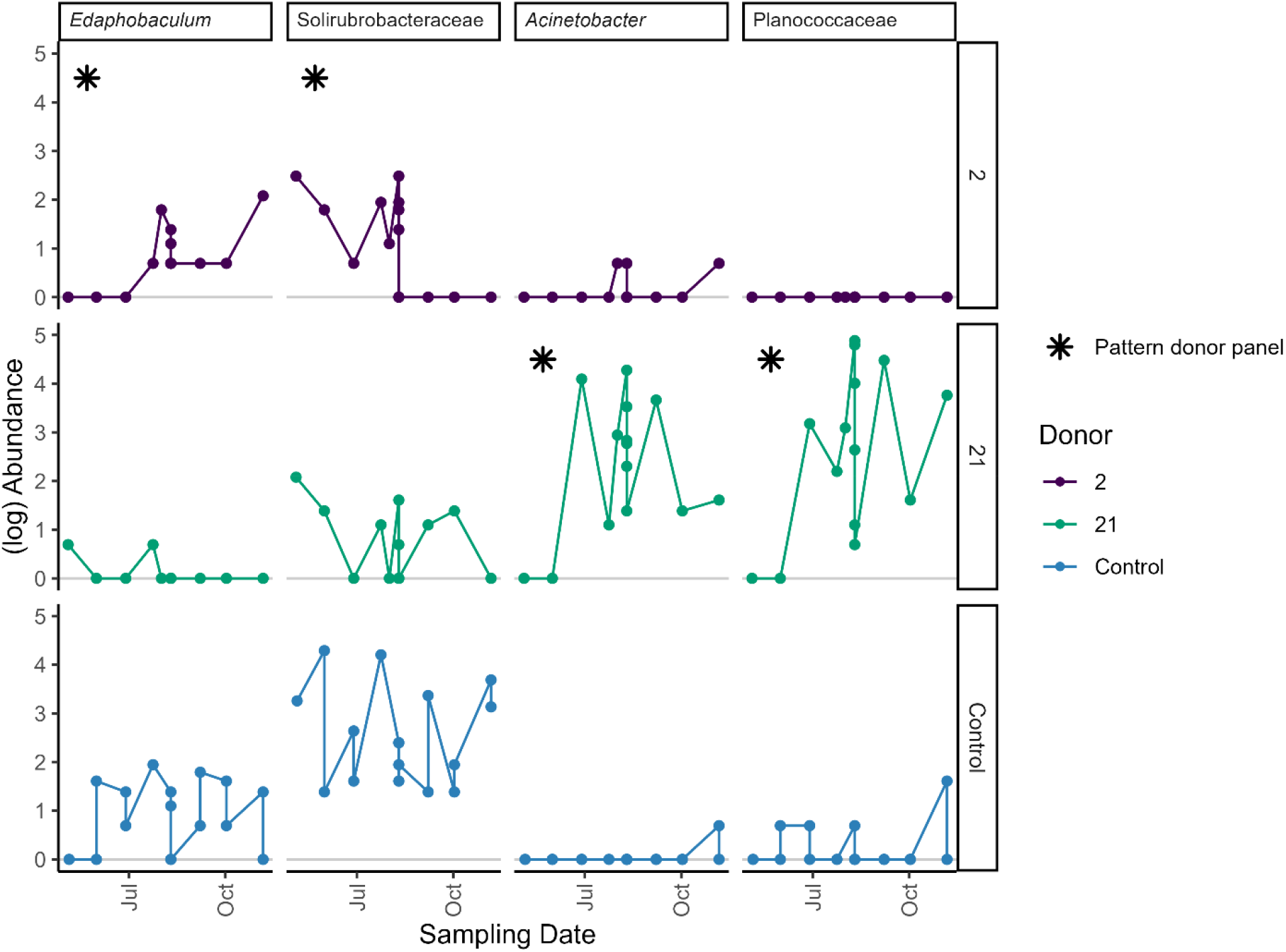
Timeseries plots of ASVs that showed appearance or disappearance patterns during decomposition. Each column represents one taxa/ASV and each donor is in one row, with the data from control sites in the third row. The panel for the donor where the appearance or disappearance pattern was observed is denoted with an asterisk. Within each panel, points are colored by donor and show log(Abundance) across sampling dates. The horizontal grey line marks log(Abundance) = 0.

## Discussion

Microbial community modelling as a method to estimate PMI has garnered increased attention in the forensic community in the past decade [15–18,58–63]. Prediction models have been developed based on soil necrobiome succession associated with human decomposition [15–18], with error rates as low as ∼3 days [18] or ∼14% of the total study period. Although these models have not yet been used to estimate PMI in a legal setting, focused research continues to add to the growing body of knowledge required to achieve error rates acceptable in the courts [64,65]. One gap in that knowledge is a lack of understanding of how spatial heterogeneity in the CDI impacts temporal relationships, which is reflected in the absence of a standardized procedure for soil sample collection. Accordingly, recently published human necrobiome studies have used variable approaches to the collection method (swab [18], core [17] or scoop [15]), collection location (anywhere within the CDI [17] or specific sites [15,16,18]) and number of samples collected per time point (single [17,18] or composite [15,16]). While within-study consistency has allowed for the identification of clear temporal trends [15–18], sampling differences have made generalization across studies difficult, and direct quantification on PMI estimates has not been formally assessed. The current study begins to address this limitation by examining spatial variation of necrobiome communities at a single timepoint and exploring how this variation impacts temporal relationships and PMI prediction.

### Spatial Variability

Observed spatial patterns in CDI soil bacterial communities were different between donors and may reflect conditions related to decomposition stage. Taxonomic diversity and richness were reduced in CDI samples from Donor 21 (sampled at PMI 257 days; 2,175 ADD), most notably in pelvic samples where soil was saturated with decomposition fluid during the advanced decay stage at the time of sampling. The sharp increase in Firmicutes abundance observed in the pelvic samples paired with the near-complete loss of other taxa is consistent with necrobiome community structures observed in other decomposition-impacted soils and suggests these selective conditions may exist unevenly within the CDI of Donor 21 [5,6,11,13,14,28,59]. In contrast, Donor 2 represented partially disarticulated bones in the dry remains stage at the time of spatial sample collection (sampled at PMI 1098 days: 10,015 ADD) and the CDI was showing signs of recovery, including plant regrowth. While these conditions are less dynamic than those associated with advanced decay, chemical and microbial signatures of decomposition have been shown to persist long after visible signs of ongoing decomposition have disappeared, with significant differences in CDI bacterial communities detected up to 10 years postmortem [7,13–15,28,66,67]. The longitudinal gradients observed in the CDI along the length of Donor 2 include increasing Proteobacteria towards the knees with higher levels of Firmicutes, Actinobacteriota and Bacteroidota at the shoulders and pelvis.

These patterns have been associated with differential exposure to decomposition products, and likely represent legacy effects of inputs from earlier stages of decomposition [13,14,19,28]. The observed shift in Acidobacteriota abundance between the left and right sides of the CDI cannot be clearly attributed to decomposition processes and may instead be a reflection of localized environmental variability that is uneven across the CDI.

Importantly, necrobiome community metrics for each donor showed no statistically significant differences between samples collected from different anatomical sites or sides of the remains, indicating that samples from any CDI location between the knees and shoulders can be expected to produce representative community data. Given the small sample size and lack of overlap in ADDs and decomposition stage between donors, there was no true biological replication across cadavers, limiting the statistical power to detect spatial differences. Although no statistically significant patterns were identified, the trends noted above may be biologically meaningful as the different patterns in community structure between donors suggests that spatial heterogeneity could vary over time, with changes in magnitude and patterns that reflect the selective pressures created by decomposition. Because of this, necrobiome community profiles from any given time point may reflect not only the temporal progression of decomposition, but also the spatial context of the CDI location from which they were collected. This has implications for PMI estimation research which relies on identifying temporal changes in necrobiome community data.

### PMI Estimation

This study identified 11 microbial taxa with statistically significant relationships to ADD in the CDI of Donor 2, indicating that a temporal signal persisted throughout the study period despite the donor remaining in the dry decomposition stage. While the base model performance varied across taxa, the presence of both positive and negative slopes suggests that the models captured successional shifts over time. Importantly, the addition of spatial samples did not alter the direction or magnitude of any model slopes, and five taxa maintained significant relationships to ADD, indicating that conditions impacting abundance of these taxa are largely consistent across CDI locations associated with dry remains. Of the models where the PMI signal was impacted (p-value became >0.05), samples from the torso-associated sites were solely responsible for the loss of significance in all but two taxa, whereas samples from the knee region exhibited more stable relationships. This spatial pattern is consistent with previous observations that areas in closer proximity to the anatomical sources of decomposition inputs experience greater disruptions to their microbial communities than areas that are more distal, where an increased persistence of indigenous soil bacteria communities was observed [14,28].

Necrobiome members in the CDI of Donor 21 did not show any statistically significant linear temporal signals. Throughout the entire sampling campaign, Donor 21 was in advanced decay, which is the period during which a large proportion of cadaver products enter the surrounding soil [1,2]. This ongoing input of decomposition products likely selected for taxa that are capable of responding to these dynamic conditions, which are accordingly unlikely to exhibit stable linear relationships. Our opportunistic approach of collecting samples from anywhere between the shoulders and pelvis may have further obscured clear linear relationships by capturing variability that reflected both spatial heterogeneity and temporal changes. Indeed, non-linear methods did successfully identify taxa with a likely PMI signal.

ASVs assigned to the genus *Acinetobacter* and family Planococcaceae appeared in the CDI of Donor 21 in the third sampling event and were present throughout the remainder of the observation period.

Members of Planococcaceae have been associated with remains throughout advanced decomposition [11,19] and may provide primary metabolic breakdown products to microbial decomposers within a necrobiome system [18]. *Acinetobacter* members have been identified as key microbial decomposers that originate from both the soil and host-associated sources and participate in a universal decomposer network that has been observed to assemble in response to vertebrate decomposition across environmental contexts [18,21]. These taxa were absent or essentially absent in the CDI of Donor 2 and control soils throughout the same period, which suggests they may only be observed within the context of active decomposition processes as opposed to background environmental conditions. This is in contrast to the two taxa that were identified in the CDI of Donor 2, which were also present in the CDI of Donor 21 and control soils. Members of the family Solirubrobacteraceae and genus *Edaphobaculum* are known soil microbes [68–70] without any documented role in vertebrate decomposition, suggesting that their presence may reflect the local soil and environmental conditions as opposed to any process related to decomposition.

## Conclusions

The necrobiome community metrics were generally consistent across CDI sites at a given timepoint, and spatial variability did not substantially impact PMI estimation. These findings combine to suggest that collecting single samples at opportunistic sites beside the remains throughout the CDI can capture PMI-informative necrobiome signals in the mid to late postmortem period, despite visibly heterogenous localized CDI conditions. This may represent a practical and robust approach for both temporal studies and the forensic investigations the resulting models aim to support, since it prioritizes preserving the decomposition environment throughout experimentation while generating predictions that are not dependent on anatomically fixed sampling locations, which may not be available in forensic recovery scenarios where remains are incomplete or have been disturbed.

