## Supplementary figures and images for "Spatial Variability in Soil Necrobiome Communities has a Negligible Effect on Postmortem Interval Estimation"

### Supplementary Figure 1

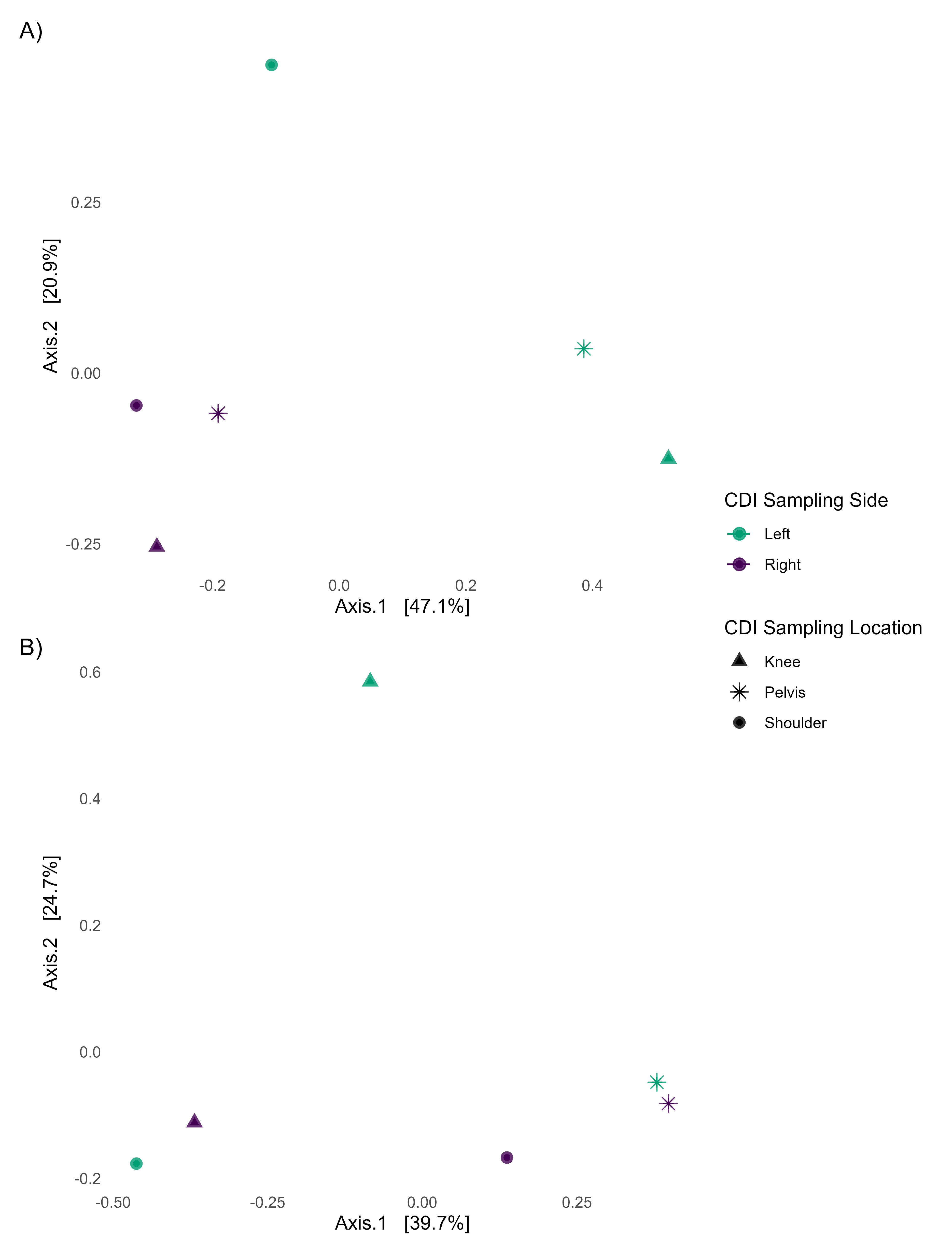
